# Genetic architecture of inter-specific and -generic grass hybrids by network analysis on multi-omics data

**DOI:** 10.1101/2022.12.23.521625

**Authors:** Elesandro Bornhofen, Dario Fè, Istvan Nagy, Ingo Lenk, Morten Greve, Thomas Didion, Christian Sig Jensen, Torben Asp, Luc Janss

## Abstract

Understanding the mechanisms underlining forage production and its biomass nutritive quality at the omics level is crucial for boosting the output of high-quality dry matter per unit of land. Despite the advent of multiple omics integration for the study of biological systems in major crops, investigations on forage species are still scarce. Therefore, this study aimed to combine multi-omics from grass hybrids by prioritizing omic features based on the reconstruction of interacting networks and assessing their relevance in explaining economically important phenotypes. Transcriptomic and NMR-based metabolomic data were used for sparse estimation via the fused graphical lasso, followed by modularity-based gene expression and metabolite-metabolite network reconstruction, node hub identification, omic-phenotype association via pairwise fitting of a multivariate genomic model, and machine learning-based prediction study. Analyses were jointly performed across two data sets composed of family pools of hybrid ryegrass (*Lolium perenne* × *L. multiflorum*) and *Festulolium loliaceum* (*L. perenne* × *Festuca pratensis*), whose phenotypes were recorded for eight traits in field trials across two European countries in 2020/21. Our results suggest substantial changes in gene co-expression and metabolite-metabolite network topologies as a result of genetic perturbation by hybridizing *L. perenne* with another species within the genus relative to across genera. However, conserved hub genes and hub metabolomic features were detected between pedigree classes, some of which were highly heritable and displayed one or more significant edges with agronomic traits in a weighted omics-phenotype network. In spite of tagging relevant biological molecules as, for example, the light-induced rice 1 (*LIR1*), hub features were not necessarily better explanatory variables for omics-assisted prediction than features stochastically sampled. The use of the graphical lasso method for network reconstruction and identification of biological targets is discussed with an emphasis on forage grass breeding.

## Background

Forage grasses cover large portions of agricultural land worldwide, efficiently converting enormous amounts of natural resources into macronutrients used primarily for feed. Their relevance can be recognized by the extent of the network of researchers and breeding organizations devoted to maximizing production efficiency. This has been largely achieved by conventional breeding techniques aiming to explore genetic variation not only within but also across species and genera over the last decades. As biotechnology surged, breeders advanced in experimenting with hybridizations across species and genera, leading to the release of successful varieties of polyploid hybrid ryegrass (L. *perenne* × *L. multiflorum*) and *Festulolium loliaceum* (*L. perenne* × *F. pratensis*), for example. As high-throughput sequencing platforms reduced genotyping costs, genomics began to play a significant role across grass breeding programs, reshaping breeding pipelines aiming at the optimization of resource allocation mainly via genome-wide selection (Keep et al. 2020;Arojju et al. 2020; Fè et al. 2015). Recently, the complex problem of predicting phenotypes and finding candidate biological molecules associated with it can also be supported not only by marker information at the DNA level but also via transcriptomics (Pignon et al. 2021) and metabolomics (Wen et al. 2014), leading to a holistic view of the phenomena controlling the expression of economically important traits.

Improving existing weaknesses of elite genetic materials or simply unlocking genetic variability for breeding exploitation are processes that benefited by leveraging hybridization across genera and species within a genus. In spite of being predominantly diploid (2n = 2x = 14), *L. perenne, L. multiflorum*, and *F. pratensis* can also be found as or induced to tetraploid states, which is essential for amphidiploid production. However, genomic instability is often reported and a shift to the ryegrass genome over generations can happen in crosses with fescues (Kopecky et al. 2017;Akiyama et al. 2012). Additionally, homeolog expression bias and expression level dominance can be observed in such allopolyploids (Glombik et al. 2021). Collectively, these phenomena may lead to distinct interactomes when hybridizations are performed across species, which can be analyzed through network reconstruction by leveraging high throughput omics data and appropriate statistical methods. Using RNA-seq data, (Hu et al. 2017) reported network topologies of allopolyploid cotton resembling more to one of the diploid species representing a progenitor besides a substantial domestication impact on the coexpression. Additional studies on expression modifications in allopolyploids remain scarce.

Adding extra layers of biological information also means increasing data dimensionality (*n* ≪ *p* problem). Reliable inferences in high dimensions require specific statistical procedures and an in-depth understanding of the underlining phenomena. Among the methods proposed for the analysis of high dimensional omics data (Bersanelli et al. 2016), the reconstruction and analysis of regulatory networks offer the possibility to prioritize omic features (Naserkheil et al. 2022), significantly reducing the searching space for downstream analyses. Organizing omic features in interacting networks can be seen as an approximation of the true existent interconnected biological system that reads the information encoded on the genome and ends with a functional organism. Reconstructed networks hold biological meaningful topological properties, for example, the presence of modules that might cluster nodes (omic features) performing specific biological functions (Li et al. 2015) and the existence of highly connected nodes. These hub nodes arise as biological networks are assumed to be scale-free, meaning that node degrees are power-law distributed (Pereira-Leal et al. 2004) and, therefore, few highly connected nodes are expected. The presence of these disproportionally connected hub nodes is an important topological property of networks as it may represent key genes/metabolites associated with biological pathways. Thus, it would be of special interest to investigate the extent to which hub omic features can be significantly linked to biomass yield and other economically important phenotypes of fodder grasses. Researchers have found hub genes affecting biomass accumulation in other families of plants, for example, in *Ulmus pumila* L.(Chen et al. 2021) and *Arabidopsis thaliana* (Liu et al. 2021). That being stated, one needs to first estimate the network to be able to explore its topological properties and this can be accomplished by leveraging graph theory and probability for modeling and representation of complex biological problems as probabilistic graphical models (Li et al. 2015).

Omics data as a graphical model is based on the estimation of conditionally independent relationships across random variables in a multivariate setting. Learning a graphic in high-dimension requires dealing with a situation where the number of unknown parameters exceeds the sample size. In this case, *ℓ*_1_-penalization has been one of the main techniques used to make sparse inference in a Gaussian Markov random field (Friedman et al. 2007), yielding a sparse structured precision matrix ∑^−1^ which, in turn, can be converted into an undirected network and further analyzed for its topological properties. This approach has been applied to the study of gene expression (Shahdoust et al. 2019; Wu et al. 2013) and metabolomic (Liu et al. 2022) data in humans, with few examples in plants(Li and Jackson 2015; Kapoor et al. 2021;e Lima et al. 2018; Bartzis et al. 2017). With a selected set of candidate features recovered from gene co-expression and metabolic networks, one can perform omic-phenotype integration. The simple correlation-based integration method of omic variables and phenotypes is widely used, with examples in maize (Zhang et al. 2019) and for the forage species *E. sibiricus* (Zheng et al. 2022). However, more robust approaches based on multivariate multi-level models have also been applied (de Steenhuijsen et al. 2016;Nantongo et al. 2021), showing better properties(Bo et al. 2014). Finding significant associations of genes and metabolites with dry matter yield and nutritive quality traits in fooder grasses could reveal potential targets for quantitative trait dissection studies, improve the omics-assisted selection of elite families, and shed light on regulatory processes of key traits. Additionally, given the fact that a large part of the above-ground biomass is harvested in forage grasses, it can be hypothesized that randomly sampled hub features are more likely to be linked to a phenotype of interest compared to, for example, grain crops.

The inherent properties of an organism’s interactome, especially the power-law distribution of interactions, give plasticity in face of random disturbances. However, interferences on hub nodes may lead to severe product alterations (Crombach and Hogeweg 2008), making them targets for genetic studies. Additionally, hub genes appear to be associated with a variety of biological processes (Tahmasebi et al. 2019; Hollender et al. 2014;Zheng et al. 2022;Hu et al. 2017) and had been mentioned as potential targets for the molecular breeding of forage species (Yan et al. 2022). In the present study, we consider the problem of reconstructing the interplay among biomolecules and narrowing down high-dimensional omics data to fewer hub features to further test their association with quantitative traits evaluated in family pools of allopolyploid grasses. Furthermore, having significantly associated hubs would confirm the relevance of these interacting biomolecules, which could be targeted for molecular biology studies and marker-assisted breeding. Our objective is, therefore, to prioritize omic features in forage grasses by sparse estimation via undirected graphical models, filter the relevant ones, and expand on their biological functions for biomass accumulation.

## Methods

### Plant material and phenotypes

Interspecific hybridization of *L. perenne* × *L. multiflorum* (hybrid ryegrass) and intergeneric crosses of *L. perenne* × *F. pratensis* (*Festulolium loliaceum*), all in tetraploidy forms (2n = 4× = 28), were performed as two connected (by *L. perenne* parents) sparse diallels in the summer of 2017 at the DLF Seeds A/S research station, Store Heddinge - Denmark. Single plants used as parents were extracted from commercial varieties of *L. perenne*, *L. multiflorum*, and *F. pratensis*. A total of 79 and 65 allotetraploid families of hybrid ryegrass and *Festulolium loliaceum*, respectively, were obtained out of several attempts. Hybrid ryegrass (referred to hereinafter as HR) families were obtained after crossing 31 *L. perenne* parents with 79 *L. multiflorum* in a sparse diallel design. For the pedigree class *F. loliaceum* (referred to hereinafter as FL), 24 *L. perenne* parents out of the 31 from the HR diallel were crossed with four *F. pratensis* parents. A sufficient quantity of seeds of F_3_ families was obtained after two rounds of multiplication. The field trials were carried out in the autumn of 2020 at two testing sites: i) Denmark (55°17′52″ N, 12°24′58″ E) and ii) the Czech Republic (49°40′59″′N, 17°58′05″’E). Families from the HR pedigree class were sown in Denmark in plots of 12.5 m^2^ with two replicates while families of the FL pedigree class were sown in the Czech Republic in plots of 6.25 m^2^, also with two replicates. At each location, entries were assigned to plots arranged in five smaller trials in a randomized complete block design with ~16 entries each. Alongside the described steps, seeds from F_2_ families were sown in a greenhouse environment in 2019 at Aarhus University, Research Center Flakkebjerg. One gram of seeds from each family was sown in pots 10 cm in diameter aiming at 120 to 150 emerging individual plants. The total above-ground biomass was harvested as one bulk per family, flash-frozen using liquid nitrogen to stop metabolism, and placed in a −80°C freezer. Frozen tissue ground into a fine powder with liquid nitrogen was used for RNA isolation and sequencing after a quality check. In addition, aliquots weighing 300 mg from ground tissue were freeze-dried for NMR-based metabolomic profiling.

We collected phenotypes for eight traits at four-time points in Denmark and three-time points in the Czech Republic across the Spring, Summer, and Autumn of the 2021 production year. The following traits were assessed: moisture-corrected dry matter yield standardized by plot size (DMY, g m^-2^), acid-detergent fiber (ADF), acid-deterged lignin (ADL), dry matter digestibility (DMDig), neutral deterged fiber (NDF), digestible NDF (NDFD), protein (Prot), and water-soluble carbohydrates (WSC). All nutritive quality traits are expressed as a percentage of DMY, except for NDFD, which is a percentage of NDF. Nutritive quality traits were obtained via a near-infrared (NIR) spectrometer onboard the plot combine harvester. Raw NIR data had previously been calibrated and is yearly updated with new wet chemistry analysis, a routine procedure in the breeding company.

### Multi-omics data

#### Gene expression via RNA sequencing

RNASeq libraries were prepared and sequenced at the Beijing Genomics Institute (BGI Hong Kong) using the BGISEQ-500RS sequencing platform technology in 100nt paired-end (PE100) mode. Paired-end reads (20 to 25M sequences per sample) were mapped to pseudo-chromosomes and scaffolds of the Lolium_2.6.1 reference genome (Nagy et al. 2022) using the splice-aware aligner HISAT2(Kim et al. 2019). Alignments were processed by StringTie (Pertea et al. 2015) for transcript reconstruction and gene expression quantification. Normalized read count values in fragments per kilobase of transcript per million (FPKM) were collected for 139,004 transcripts annotated on the Lolium_2.6.1 reference genome. A filter was applied to the expression profile matrix to get rid of transcripts with expression values very low/equal to zero. The threshold for transcription was set to 0.5 median FPKM across all samples, yielding the final filtered gene expression matrix with 18,499 transcripts.

#### RNASeq-based genetic variants

Variant calling was performed from RNA-seq merged BAM-format alignments using the Bayesian genetic variant detector Freebayes(Garrison and Marth 2012). The initial single-nucleotide polymorphism (SNP) calling resulted in 1,689,206 variants. After retaining only biallelic markers, we filtered variants by the following criteria: i) a maximum missing proportion of 50% at each locus, ii) a minimum mapping quality of 20, iii) a minimum read coverage of five reads per variant position, and iv) minor allele frequency (MAF) greater than 0.05. The final set of SNPs comprises 89,862 variants that were used for downstream analyses.

#### NMR-based metabolomic data

The metabolomic profiling by proton nuclear magnetic resonance spectroscopy (1H-NMR) was carried out at the Natural Products Laboratory (The Netherlands). Following the sample preparation and spectra acquisition with a 600 MHz Bruker AVANCE III spectrometer (Bruker BioSpin GmbH, Germany), the raw NMR data were processed using the software package NMRProcFlow (Jacob et al. 2017). After chemical shift calibration and normalization, metabolomic fingerprinting yielded a total of 556 bins with non-zero intensities (referred to hereafter as NMR variables) for 144 plant samples by applying an adaptive Intelligent Binning [AI-Binning, (Meyer et al. 2008)] algorithm. A tab-separated file with samples on rows, NMR variables on columns, and cell-wise intensity values was generated for downstream analysis.

### Statistical analysis

Prior exploratory analysis revealed considerable differences between the omics data from the HR class compared to FL class samples. Therefore, downstream analyses were performed considering each of the two classes as distinct but related across layers of omics data. Additionally, this decision was supported by the fact that phenotypes were assessed in different locations, lacking connectedness. Later, these data sets were merged for an omic-assisted prediction study.

#### Allele frequency-based genomic kernel

The genomic relationship matrix (GRM), which gives the realized genetic similarities among any pair of individuals, was computed for SNP data sets of sizes *p* × *n* equal to 85,283 × 79 for the HR and 75,299 × 65 for FL data sets after individually re-filtering by MAF, depth, and missing rate using the same thresholds as described before. The GRM was then used for downstream omics feature corrections due to population stratification and multivariate mixed model analysis. The GRM based on pooled DNA was calculated using (VanRaden 2008) method 2 adapted to use allele frequencies instead of discrete genotype calls. First, a column-centered matrix **M** was computed as 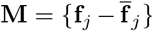, with *j* indexing SNP markers, **f**_*j*_ representing a vector of alternative allele frequencies for SNP j, and {…} represents a matrix built up with column vectors. The matrix **G** can then be obtained as shown in Eq. 1.

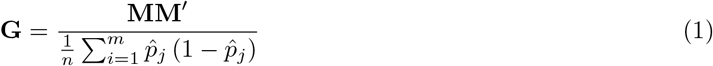

where *n* is the ploidy of the breeding material, *m* is the number of markers, and 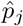 represents the frequency at *j*th locus simply obtained by taking the column means of the **M** matrix. As outbred full-sib **F**_2_ families of tetraploid plants, the genotype of a family can be described as octoploid (Ashraf et al. 2014). Therefore, the realized relatedness is obtained by scaling the plain genomic relationship matrix from the cross product of **M** by the expected SNP variances, yielding a kernel that is analogous to the traditional numerator relationship matrix, also known as the **A** matrix. Finally, a diagonal correction was applied to **G** considering ploidy number and coverage depth as proposed by (Cericola et al. 2018).

#### Adjustment for population stratification

The impute file for the analysis of gene expression data consisted of two subsets of 4,767 features times the number of samples of each pedigree class. The reduced set of genes was obtained after further filtering out transcripts with more than 50% of samples having zero reads and retaining positions with at least 10 or more samples having 10 or more reads. Additionally, a filter on the expressional variance of non-zero elements was performed, selecting features ranked in the top 50th percentile as the variation for genes in the bottom may be largely due to non-biological noise. Finally, we retained only features common to both data sets followed by the addition of a pseudo count to the expression matrix, which was subsequently log(2)-transformed [log_2_(x+1)]. The input file for the analysis of NMR data consisted of two subsets of 556 NMR variables for each pedigree class. NMR features were mean-centered and variable intensities were addressed via Pareto scaling, which uses the square root of the standard deviation to reduce the relative importance of high-variance features across the spectrum without much disturbance to the data structure.

Population stratification was detected in an unsupervised manner via the multivariate statistical technique of principal component analysis and corrected via regression modeling. We empirically retained coordinates of the top 10 eigenvectors of each *k* pedigree class to regress out population stratification as well as possible batch effects among samples. Therefore, the transcriptomic and metabolomic data sets were feature-wise corrected by incorporating principal component scores in the linear model of the form described in Eq. 2.

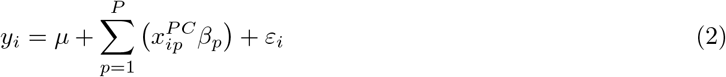

where, *y_i_* represents the response variable *i* (omic feature); 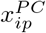 is the entry-specific coordinates of the *p*th principal component, with *p* =1…*P* where *P* is equal to 10, *β_p_* is the fixed regression coefficients adjusting for population stratification, and *ε_i_* is the residual which was retained to reconstruct the full corrected omics data sets for network estimation.

#### Joint graphical lasso analysis for inverse covariance estimation

A joint graphical lasso (JGL) method was used for estimation in a scenario of double-related Gaussian graphical models. The two-class problem of high dimensional features was present in the data set due to the available inter-species/genus crosses. One can expect similar graphical models between the two classes as parents were shared among crosses between them, but also some nuances once the involved species have substantial differences regarding phenotypic traits. Therefore, the joint graphical lasso proposed by (Danaher et al. 2014) can handle this situation by estimating two graphical models, one for each pedigree class, and borrowing information across classes. For each pedigree class *k* (*k* = 1, 2), let a data matrix **X**^(*k*)^ represent column-centered data with p omic features, and **X**^(*k*)^ ~ *N* (*μ*^(*k*^, **Σ**^(*k*)^), where **Σ**^(*k*)^ is a positive definite *p* × *p* covariance matrix of the omic features. The inverse of **Σ**^(*k*)^ is the precision matrix **Θ**^(*k*)^ representing the network structure of omic features. By applying an *ℓ*_1_-penalty on **Θ**^(*k*)^ the network is made sparse, where elements will be 0 for conditionally independent pairs of features given the remaining variables. The sparsity condition allows learning graphics even in small sample sizes. The fused graphical lasso formulation in which **Θ**^(*k*)^ are estimated by maximizing the penalized form of the likelihood function for the two classes is shown in Eq. 3.

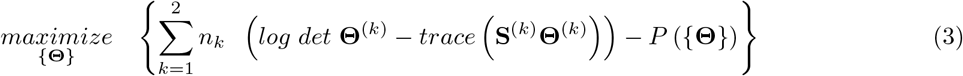

where *P* ({**Θ**}) is as follows:

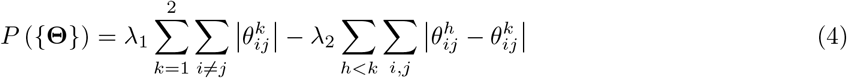

here, **S**^(*k*)^ is the empirical covariance matrix of omics features calculated as **S**^(*k*)^ = *n*^-1^**X**^(*k*)^**X**^(*k*)*T*^. The optimization problem is here solved by the alternating direction method of multipliers (ADMM) algorithm. The solution to the problem of *n* ≪ *p* in the joint graphical lasso model is based on a penalized log-likelihood approach. In addition, as can be seen in Eq. 4, running JGL requires tuning two nonnegative parameters (λ_i_ and λ_2_). The λ_1_ penalty controls the degree of sparsity while λ_2_ determines network similarity. If λ_2_ is zero (i.e., no penalty is imposed) then **Θ**^(*k*)^ are independent and no information is shared between them. To select the proper hyperparameters, we used a goodness-of-fit approach where a grid search was performed to select values that minimize the Bayesian information criterion [BIC] (Schwarz 1978) specified in Eq.5 (Augugliaro et al. 2016), yielding values that balance model likelihood and complexity.

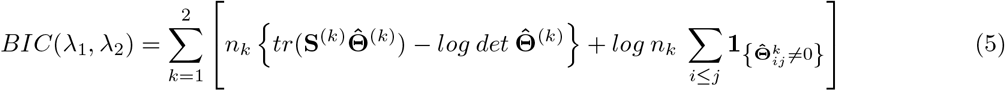

In order to reduce the computational burden, a dense search was performed over λ_1_ for each fixed value of λ_2_ and a quick search for the former parameter for each fixed value of λ_1_ as suggested by (Danaher et al. 2014). For the metabolomic data set, a uniform log spaced grid starting from 0.01 to 20 with a size equal to 30 was defined for λ_1_ whereas a simple sequence equally spaced from 0 to 0.5 (size of 15) was defined for λ_2_. The same grid search space was defined for transcriptomic data, however, smaller sizes of 15 for λ_1_ and 10 for λ_2_ were specified. After selecting the proper hyperparameter values, we run JGL for each omics data set producing four precision matrices **Θ**^(*k*)^. From these matrices, one can compute the partial correlation between pairs of dependent features as 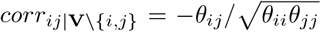. The joint graphical lasso method implemented in the R package JGL (Danaher et al. 2014) was used for network estimation.

#### Network reconstruction, candidate modules, and hub identification

Network analyses aiming for complexity reduction were performed in order to prioritize candidate genes and metabolomic features for further integration with phenotypes of interest. Initially, each precision matrix **Θ**^(*k*)^ was converted into a symmetric (graph is undirected) 0-1 matrix of dimensions equal to *p* × *p*, referred to as the adjacency matrix **A**^(*k*)^ for each *k* data set following the definition:

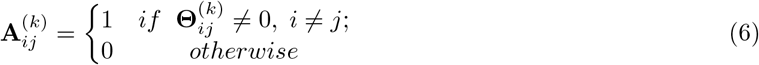

Four adjacency matrices **A** were obtained and from them, we created graphic objects using the R package igraph (Csardi and Nepusz 2006). Initially, a graph is denoted as *G* = (*V, E*) in which each node *ν* ∈ *V* represents a biomolecule in this study, whereas each edge *e* = (*ν_i_*, *ν_j_*) ∈ *E* refers to the interaction between pairs of nodes *ν_i_* and *ν_j_*. Each graph was organized in modules (communities) via a multi-level modularity optimization algorithm (Blondel et al. 2008), forcing highly connected edges to cluster in modules that are sparsely connected among them. In other words, more edges occur within identified modules than the quantity expected at random. The community structure is essential for finding hub nodes that are more likely to be involved in different biological processes.

Hub features were identified intramodule via maximum Kleinberg’s hub centrality score, which is the principal eigenvector of **A**^(*k*)^ · (**A**^(*k*)*T*^ (Kleinberg 1999). By using the hub scores, one can identify the most influential features in the network and explore the biological function of these interacting biomolecules. Therefore, we selected the top five hub features per module and kept only those intersecting across data sets to maximize the probability of selecting true/conserved hubs of genes and metabolites.

#### REML variance components and heritability

Single omic features were analyzed by fitting a linear mixed model of the form: **y** = **1***μ* + **Zu** + **e**, where **y** is the response vector (normalized gene expression values or total area of the bin from the bucketed NMR spectrum), **1** is a vector of ones linking observations to the constant *μ*, 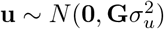, and 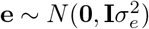 are vectors of the random additive genetic with covariance structure **G** (Equation 1) and independent (identity matrix **I** as covariance structure) residual effects, respectively. **Z** is the design matrix assigning observations of omic features to the respective F_2_ family. The genomic heritability was calculated as 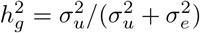, where 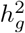 measures the proportion of the variance attributed to allele substitution effects captured by the genomewide markers relative to the total variance.

Phenotypic variance within location was partitioned into the terms defined by the linear mixed model displayed in Equation 7:

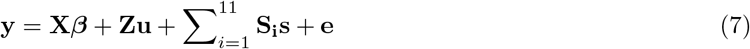

where, **y**, *β*, **u**, **s**, and **e** represent the vectors of the response variable, fixed trial-block effect, random additive genetic effect following 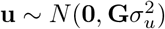, random spatial effect following 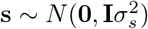, and random residual effect assumed 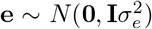, respectively. Matrices **G** and **I** are as defined before. Design matrices **X**, **Z**, and **S** link observations of the response variable to the specific model effect. The spatial effect is a sliding window accounting for 10 neighboring plots in addition to the target experimental unit and works by scanning the field for spatial variation not accounted for by the prior trial design. Genomic heritability was calculated as: 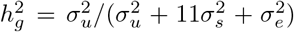. Variance components and heritabilities for eight phenotypic traits can be found in the supplemental Table S1. Finally, the parameter 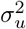 was multiplied by the average diagonal of the GRM in both heritability equations presented before.

#### Phenotypes and omics integration via pairwise fitting of mixed models

The raw phenotypic data were analyzed alongside hub omic features in a multitrait genome-wide fashion via linear mixed models to investigate pair-wise additive genetic correlations. The bivariate model (Eq.8 and 9) was fitted *lm* times, combining *l* hub nodes and *m* phenotypic traits, for each data set, yielding correlations used to describe the existence of a significant association between the concentration of selected biological molecules and economically important phenotypes.

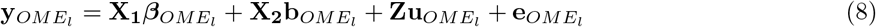

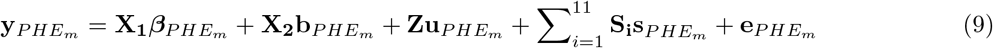

where **y***_OME_l__* and **y***_PHE_m__* are vectors of expression/intensities of hub omic features and records of phenotypic traits, respectively; *β_OME_l__* contains the fixed general mean effect while *β_PHE_m__* also contains the fixed effect of block within trial; vectors **b***_OME_l__* and **b***_PHE_m__* contains fixed regression coefficients estimated by regressing response variables on principal components’ dimensional scores calculated from the genomic kernel; **u***_OME_l__* and **u***_PHE_m__* are vectors of families’ additive genetic effect; **s***_PHE_m__* is the vector of random spatial effect with 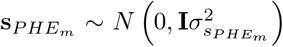; and **e***_OME_l__* and **e***_PHE_m__* are vectors of random residuals for expression/intensity of hub omic feature *l* and phenotypic trait m, respectively. For incidence matrices **X** linking fixed effects to response variables, the general mean was the only fixed effect for submodel 8, thus **X**_1_ = **1**. Matrices **X**_2_ contain scores of the top three principal components computed from the **G** matrix (Eq. 1) instead of 1’s and 0’s, aiming at further accounting for population structure to avoid false-positive associations. The selection of the appropriate number of PC’s followed an empirical evaluation of the changes in response variables’ heritabilities as they were added. The matrix **Z** is the corresponding incidence matrix of additive family effects. Finally, the series of matrices **S**_i_ link the random spatial effect to the surrounding plots and work as a sliding window (cross-shaped format) mapping the field for microenvironmental variations missed by the blocking design. The joint covariance structure of the remaining random terms was assumed as follows:

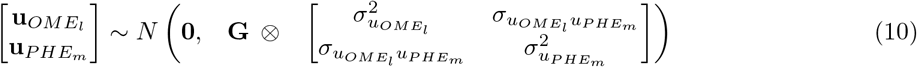

and

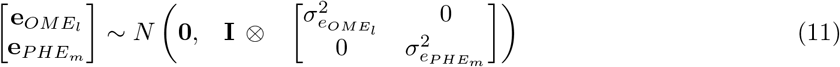

where **I** represents an identity matrix and ⊗ is the Kronecker product. Besides the scores of the first three principal components, here **G** also accounts for the whole-genomic relationship structure of the population. Covariances between response vectors were set to non-existent for residual genetic and error random effects. For hypothesis testing, we also ran a constrained version of the bivariate model, setting the additive genetic covariance between submodels 8 and 9 (Eq. 10) to zero (*σ_u_OME_m___ u_PHE_m__* = *σ_u_OME_l___ u_PHE_m__* = 0). The significance of the additive genetic correlations was tested by comparing the constrained and unconstrained models via a one-tailed log-likelihood ratio test (LRT) with 0.5 degrees of freedom (Gilmour et al. 2015; Self and Liang 1987). Multiple testing correction was performed for coefficients across traits within omic features via Benjamini-Hochberg false discovery rate (FDR)(Benjamini and Hochberg 1995) procedure at alpha equals 0.05 aiming to control for type I error.

The *lm* additive genetic correlations estimated by fitting the full bivariate model for each data set were retained along with the p-values and FDR-based significant associations and used for constructing the omics-phenotype weighted network graph. A visualization of the network was produced using the software Cytoscape 3.9.1 (Shannon et al. 2003), weighing edges by the magnitude of the trait-omic associations.

#### Gene ontology enrichment analysis

Transcript protein sequences were subjected to local InterPro analysis using InterProScan v5.28-67.0 (Jones et al. 2014). Predictive information concerning conserved protein domains, signal peptides, transmembrane domains, and gene ontology (GO) data was acquired from 14 member databases of InterPro. Per transcript, non-redundant GO information was collected from InterPro outputs using custom scripts. GO-term enrichment analysis was carried out using the Python library GOATOOLS (Klopfenstein et al. 2018) by intersecting the GO-term list of the full perennial ryegrass transcriptome, the GO-term subset of expressed genes, and the GO-term lists of filtered transcript sets (study lists). Significant enrichment was declared via Fisher Exact Test, corrected for false discovery rate (Benjamini and Hochberg 1995).

#### Omics-assisted prediction

Starting from the centered **M** matrix of SNP markers defined before, missing allele frequencies were imputed by chained random forest. This method was selected after comparing the ability in predicting missing allele frequencies against the weighted K-nearest neighbors (KNN) method via cross-validation. The imputations were performed for each pedigree class separately using the R package missRanger (Mayer 2021). The missRanger function ran using the arguments *num.trees* equal to 100, *sample.fraction* equal to 0.1, *max.depth* of 6, and extratrees for the *splitrule* argument. The imputation was performed by looping over one chromosome at a time within clusters of SNPs created by running a complete-linkage clustering algorithm with *k* = 30 as the desired number of groups.

We used the best linear unbiased estimator (BLUE) of entries as response variables in the prediction study. The adjusted phenotypes were obtained by rearranging the terms and refitting the submodel in Eq. 9 with families as a fixed effect and no PC scores were included. BLUEs within locations were mean-centered to remove differential environmental effects followed by the merging of phenotypes and predictors from HR and FL data sets. The unsupervised machine learning algorithm random forest was used as the engine for the prediction study. Models were fitted using the ‘ranger’ R package (Wright and Ziegler 2017) with the hyperparameters minimum node size and a number of randomly drawn candidate features set to five and 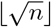, respectively, where n is the number of variables. Therefore, the random forest model was fitted on the combined data sets, setting the number of decision trees to 2,000. Training out-of-the-bag accuracy (OOB accuracy) was reported as a performance metric. Finally, variable importance was computed via permutation.

Three prediction scenarios were studied. First, we selected a subset of SNPs tagging common hub genes across data sets, the common hub genes, and the common hub NMR variables as three sets of regressors. The second scenario consisted of stochastically sampling 20× sets of 30 genes (then SNPs within these genes) and 32 NMR variables aiming to compare the prediction power contained in hub nodes with randomly sampled features. In the last scenario, we used all common SNPs, genes, and NMR variables as regressors. Besides comparing prediction accuracy with the previous scenarios, here we can assess a common prediction task where the goal is to evaluate the closeness of predicted and observed values using all available predictor variables.

#### Statistical computing and data visualization

Large-scale computations were performed in the GenomeDK high-performance computing facility located at Aarhus University, Denmark. Mixed model analyses were fitted using DMU package version 6 (Madsen and Jensen 2013). Modular network visualizations were produced using the R package NetBioV (Tripathi et al. 2014) with the Fruchterman-Reingold layout algorithm to arrange nodes in each module. Finally, miscellaneous plots wore drawn employing the ggplot2 R package (Wickham 2016).

## Results

### Genetic similarity among family pools and omics heritability

We constructed a genomic relationship matrix (GRM) for the *L. perenne* × *L. multiflorum* (hybrid ryegrass; HR pedigree class) using 85,283 SNPs and a GRM for the intergeneric crosses of *L. perenne* × *F. pratensis* (*Festulolium loliaceum*; FL pedigree class) using 75,299 SNPs (Figure 1 A and B, respectively). The average genomic relationship was close to zero as expected due to the centering of allele frequencies in both data sets (−0.0178 and −0.024 for hybrid ryegrass and *F. loliaceum*, respectively) but with substantially more variation found in the FL data set (off-diagonal standard deviation equal to 0.21 compared to 0.15 in the HR class). In addition, GRM heatmaps are substantially populated with negative relationships, meaning that many pairs of individuals were less related than the average genomic relationship. Also, the GRMs revealed biparental combinations that substantially deviated from the expected offspring composition of bi-parental crosses of single-plant parents, suggested by the presence of blocks of high genomic relationships (>1.0) among families, especially for the FL data set (Figure 1B). For instance, the 4×4 block on the top-left side of Figure 1B holds highly related families that share the same pollen receptor parent crossed with different *F. pratensis* genotypes. As the diallel design was not accounted for, downstream analyses were performed controlling for population stratification due to replicated parents in the crossing scheme using principal component (PC) scores as covariates. The first 10 PCs of the GRM matrices explained a cumulative percentage of variation equal to 75% and 82% for HR and FL data sets, respectively. Additionally, adjusted means on the right-hand side of Figure 1 reveal blocks of families with similar trait-specific performance as they were hierarchically clustered by IBS-based measurement of relatedness.

**Figure 1:**
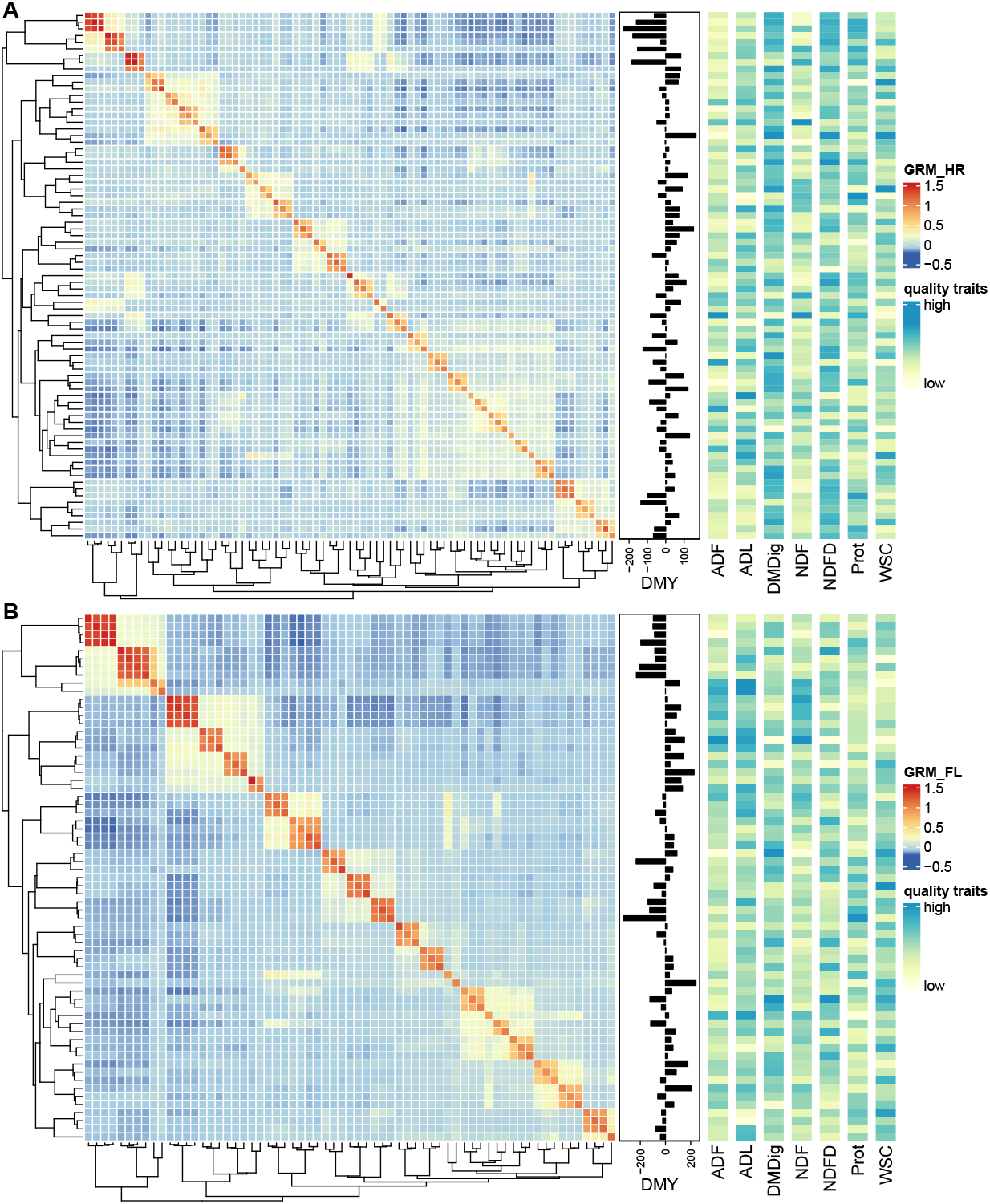
Allele frequency-based genomic relationship matrix (GRM) for 79 families of hybrid ryegrass [HR] (A) and 65 families of *Festulolium loliaceum* [FL] (B). Heatmap depictions of GRMs are annotated with best linear unbiased estimators (BLUEs) for dry matter yield (DMY) and each of the seven nutritive quality traits: ADF - acid detergent fiber; ADL - acid detergent lignin; DMDig - digestible dry matter; NDF: -neutral detergent fiber; NDFD - digestible NDF; Prot - protein; and WSC - water-soluble carbohydrates. Partially surrounding dendrograms were produced using Euclidean as the distance measure and the agglomerative complete-linkage method to build the hierarchy of clusters.

The GRMs displayed in Figure 1 were also used in a linear mixed model to estimate the genomic heritability of NMR variables and gene expression entities. The density plots of the heritabilities for both pedigree classes are displayed in Figure 2. For the HR class, median heritabilities of 0.047 and 0.122 with an interquartile range (IQR) of 0.177 and 0.311 were observed for NMR variables and gene expression, respectively. For the FL class, we observed median heritability of 0.162 and 0.165 with IQR of 0.273 and 0.295 for NMR variables and gene expression, respectively. Distributions are positively skewed and a higher quantity of high heritable variables can be detected for gene expression data in comparison to NMR variables. Additionally, the figure suggests a slightly higher proportion of more heritable features measured on samples from the FL class, especially for metabolomic data. Finally, subfigures 5B and 5C reveal the similarity in heritability between pedigree classes according to the spectrum and genomic position, respectively. Overall, there is a high correspondence between classes for regions displaying high and low heritability.

**Figure 2:**
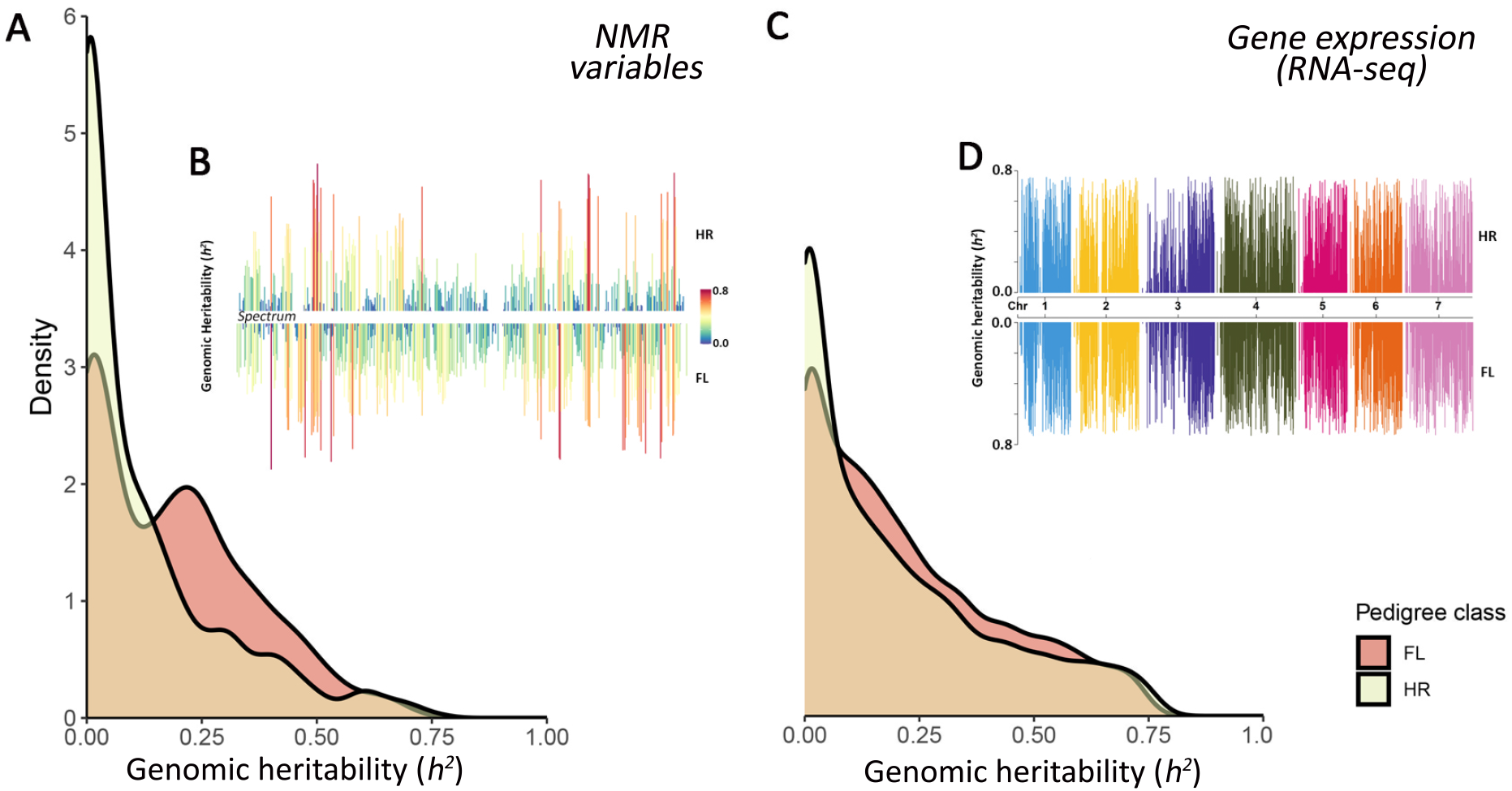
Density plots displaying the SNP-based genomic heritability distribution of NMR variables (A) and gene expression (C) from family pools of two pedigree classes (HR: hybrid ryegrass and FL: *Festulolium loliaceum*). The genomic heritability of NMR variables is displayed along the spectrum for both pedigree classes in B and subfigure D displays the genomic heritability of gene expression data along genomic position across chromosomes also for both pedigree classes.

**Figure 3:**
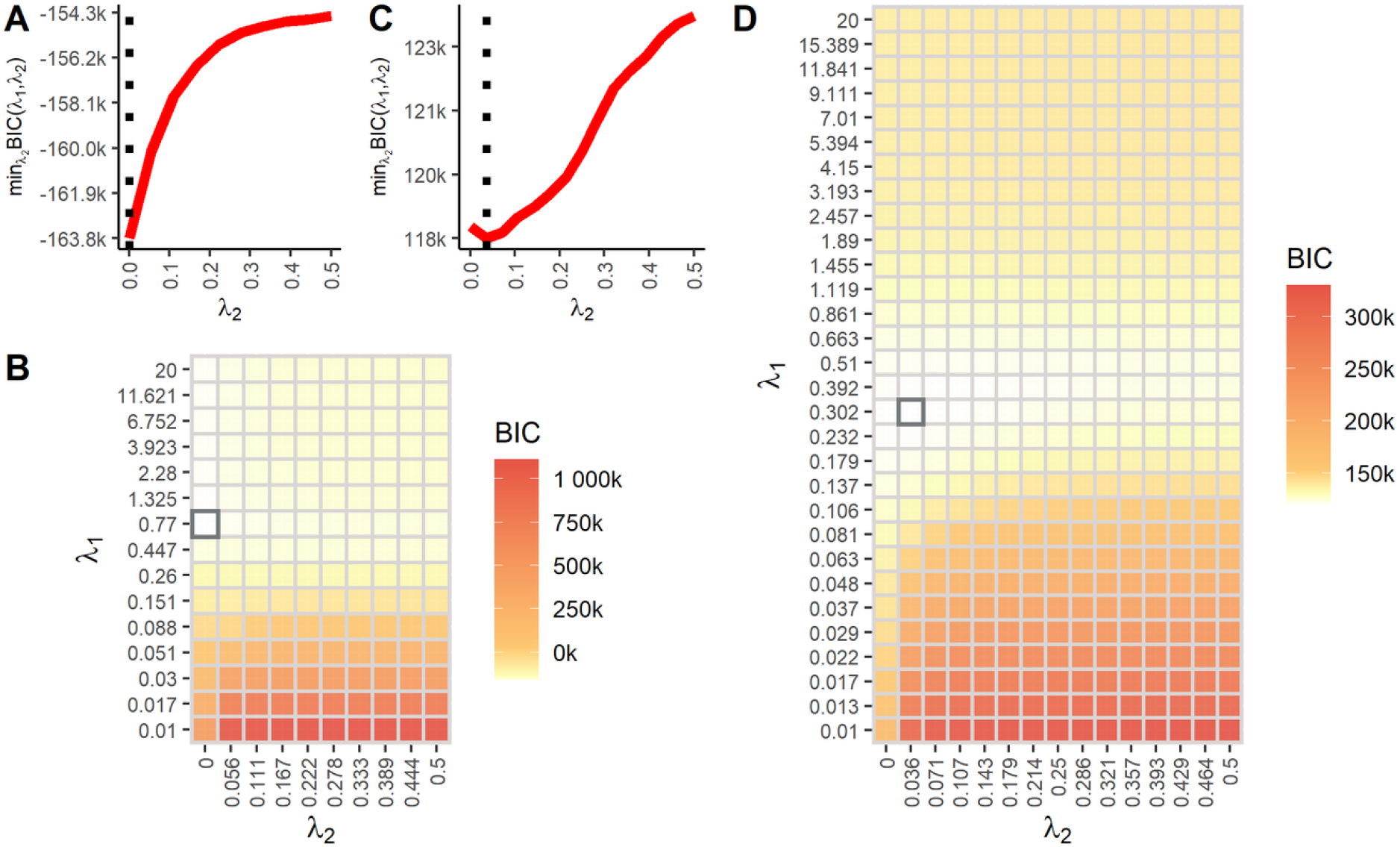
Grid searching of hyperparameters for graphical lasso model selection with *ℓ*_1_ regularization. A and C shows the Bayesian information criterion (BIC) as a function of the second (λ_2_) penalty for transcriptomic and metabolomic data sets, respectively. B and D are heatmaps displaying the complete grid search for the values of the tuning parameters λ_1_ and λ_2_ that minimize BIC, yielding parsimonious models for transcriptomic and metabolomics data sets, respectively.

**Figure 4:**
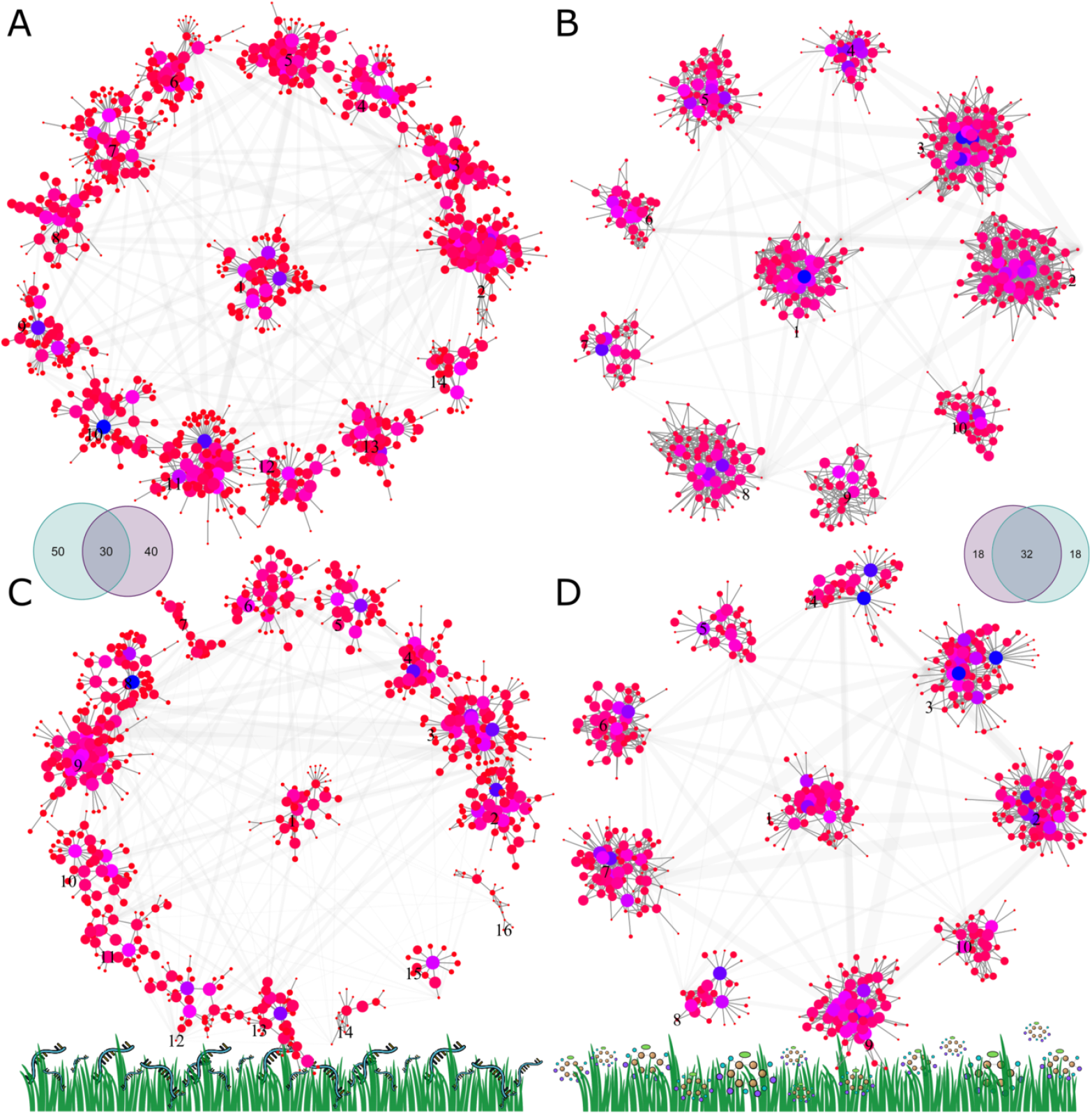
Abstract modularity view of the gene co-expression and metabolite networks constructed from gene expression data (A and C) and nuclear magnetic resonance (NMR) spectroscopy (B and D) for two data sets (A and B is the HR data set; C and D is the FL data set). Both node color and size reflect the hub score, i.e., the principal eigenvector of **A** · *t* (**A**) matrix operation, where **A** is the adjacency matrix of each graph. The color range goes from red for low-degree nodes to blue for highly connected ones. Edges between modules were collapsed and the width refers to the number of connections shared between any two modules. Venn diagrams show the overlap among sets of top hub features from each data set.

**Figure 5:**
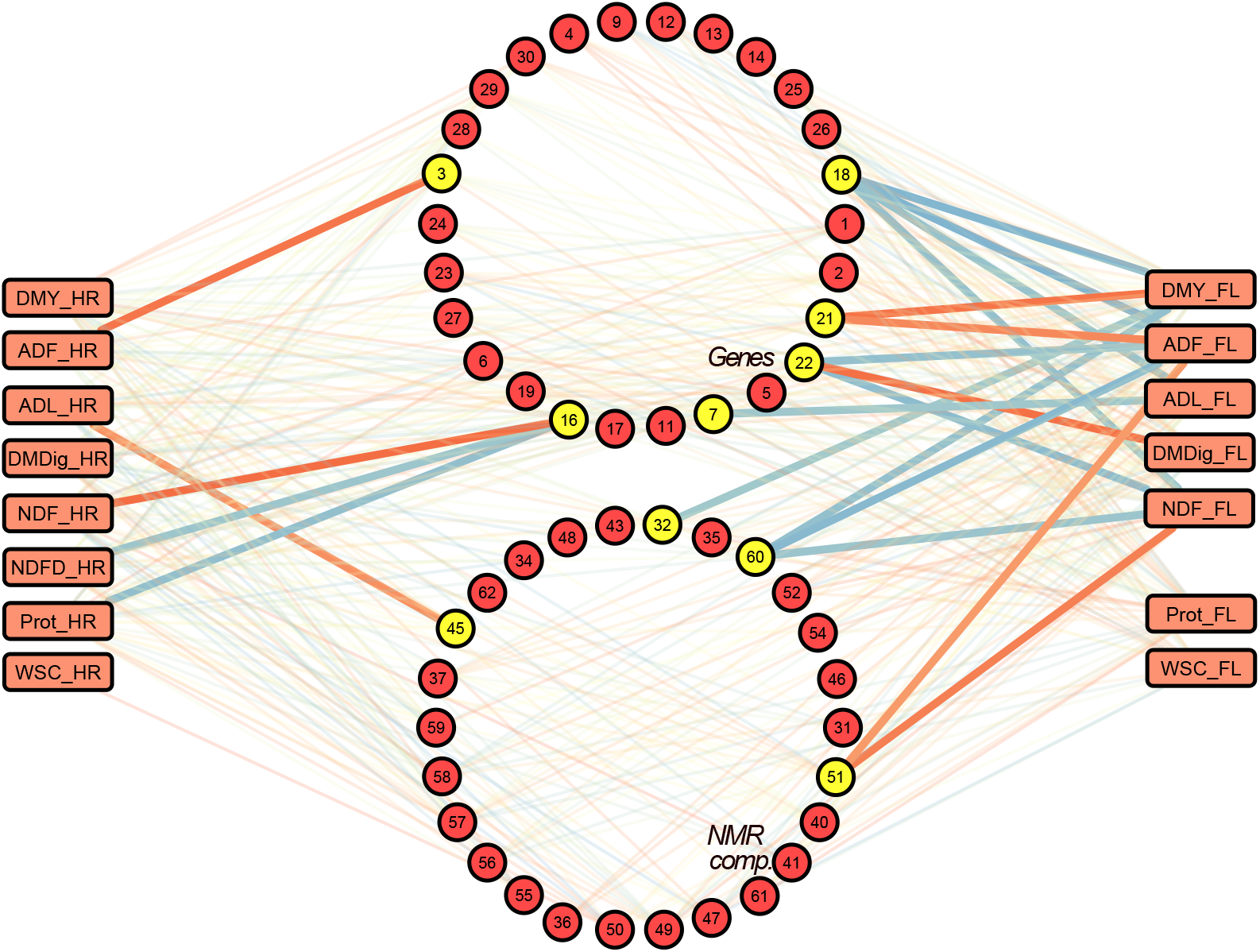
Weighted network linking hub omic features to phenotypes collected from family pools of two fodder grass pedigree classes (HR [hybrid ryegrass] data set on the left side and FL [*F. loliaceum*] data set on the right side). Edges represent the additive genetic correlation between omic features and traits and were built by the pair-wise fitting of a multivariate genomic model. Stronger edges in a gradient from red (negative) to blue (positive) colors represent false discovery rate corrected significant correlations at alpha 0.05. Highlighted omic nodes show at least one significant edge. DMY - dry matter yield; ADF - acid detergent fiber; ADL - acid detergent lignin; DMDig - digestible dry matter; NDF: -neutral detergent fiber; NDFD - digestible NDF; Prot - protein; and WSC - water-soluble carbohydrates.

### Hyperparameter tuning of joint graphical lasso

The search for the appropriate values of λ_1_ and λ_2_ that returned the smallest Bayesian information criterion (BIC) was computationally intensive as the model was fitted for all combinations of the penalties defined in the grid search, requiring several days of CPU time for joint graphical lasso (JGL) model of transcriptomic data but only using few wall time hours by taking advantage of multi-core processing. A total of 939 connected nodes were estimated for gene expression. Within data sets, four sparse subnetworks and 4,038 edges were obtained for HR whereas five sparse subnetworks and 2,182 edges were identified for the FL class given the tunning parameters selected via BIC (Figure 3). Additionally, 462 edges were found to be shared by the two pedigree crossing classes. For the next omic layer, all 556 nodes (NMR variables) were connected, one sparse network on each pedigree class was estimated, 7,757 and 4,789 edges were available for HR and FL data sets, respectivelly, and 2.371 common interactions shared by all classes.

For the gene expression data, λ_2_ was optimized at λ_2_ = 0. This implies different networks for each pedigree class with a different arrangement of non-zero positions for the gene expression data. On the other hand, for NMR data, the best combination of λ_1_ and λ_2_ that minimized the BIC found a small non-zero value for λ_2_, implying a small level of similarity on the sparsity pattern across precision matrices for NMR data. Overall and across omic layers, the hybridization process generated substantial differences between pedigree classes and it seems to be better captured at the gene expression level.

### Exploring lasso penalized precision matrices and network topologies

We detected 14 candidate modules for gene expression and 10 modules for metabolomic for the HR class (Figure 4). In the FL data sets, it was estimated 16 modules for gene expression and also 10 modules for metabolomics data. The modularity view of the gene-to-gene and metabolite-to-metabolite networks reveals the power-law distribution of node connections, where few vertices are highly connected whereas the majority has only one or few connecting edges. The organization of network structure based on modularity optimization allowed for the selection of intramodular hub nodes that are more likely to be involved in different biological pathways. Out of 70 hubs extracted from HR transcriptomic data (Figure 4A) and 80 from FL transcriptomic data (Figure 4C), 30 genes (hubs) were conserved. These high-degree genes are located across all seven chromosomes, varying from two hubs on chromosome three up to 10 on chromosome two. Also, the degree of the hub gene set ranged from 34 to 182 edges. For metabolomic data, we found 32 conserved hub nodes (Figure 4, B and D), all localized in one half of the NMR spectrum and with degrees ranging from 52 to 357 edges.

### Integrative omics

The pairwise fitting of the multivariate genomic model revealed 21 significant edges between traits and omic hub features after FDR correction (Figure 5). The multi-trait model was fitted 496 times but failed to converge in 54 cases, possibly due to the variance component being close to zero. Therefore, five traits displayed at least one significant edge with hub features in both pedigree classes. More edges can be seen on the left side of the omics-phenotype network relative to the right side, which can be explained by the higher heritability across traits in the FL data set (Supplemental Table S1) as well as overall higher heritability of genomic features (Figure 2). Additionally, significant connections were found for six out of 30 hub genes and four out of 32 hub NMR variables. Three (hubs 16, 18, and 21) out of the six genes are located distantly apart on chromosome four whereas the remaining hubs 3, 7, and 22 are located on chromosomes one, two, and five, respectively. Genomic heritabilities of hubs displaying significant edges were considerably higher compared to the full feature space, with median *h*^2^ twice as large. A closer look reveals a consistent pattern regarding the direction of the associations. Hub features positively or negatively associated with fiber content traits are also positively or negatively associated, respectively, to dry matter yield. The same holds true for protein content and digestibility traits, where associated hub features are inversely connected to fiber content. Additionally, the majority of hubs associated with phenotypes have more than one significant edge computed from independent analysis and, therefore, confirms the reliability of the estimated omics-phenotype network. We also fitted hub features as covariates in submodel 9 and computed the z-scores and associated p-values, which overall confirmed the results displayed in Figure 5 (data not shown). Finally, no hub feature had significant edges with traits from both pedigree classes, which can suggest steady genetic differences between classes and/or a lack of power to detect these shared genomic-based associations.

Gene-set enrichment analysis revealed four gene ontology (GO) terms enriched in the set of 30 hub genes displayed in Figure 5. Overrepresented GO terms were GO:0019438 (aromatic compound biosynthetic process), GO:0018130 (heterocycle biosynthetic process), GO:1901362 (organic cyclic compound biosynthetic process), and GO:0044271 (cellular nitrogen compound biosynthetic process). Bivariate mixed model analysis revealed significant genetic correlations between the expression of gene hubs 18 and 21 and dry matter yield. While hub gene 18 codes for the *atpF* gene (synthase subunit b, chloroplastic) and is associated with energy production (GO:0015986 - proton motive force-driven ATP synthesis), the blast of biological sequences revealed a putative unclassified retrotransposon protein originating from hub gene 21.

### Omics-assisted predictions

Using gene expression data as an independent variable performed similarly to SNP-based marker predictions, except for digestibility, protein, and neutral detergent fiber (Figure 6). Despite the overall poor prediction performance across traits obtained when using NMR features as independent variables, the information contained in this omic layer is useful for protein content prediction, with correlations above 0.4. Prediction accuracy using only hub genes was compared with a second scenario where samples of the same size were drawn from the whole predictor space aiming to check whether hub features carry asymmetrically more (or less) information for prediction purposes. Overall, hub NMR variables appear to be more predictive of nutritive quality traits than random samples of metabolomic features. On the other hand, results suggest a weaker relationship between observed and predicted quality parameters using hub genes as regressors. Finally, using the whole set of available predictors yields predominantly higher accuracies across traits.

**Figure 6:**
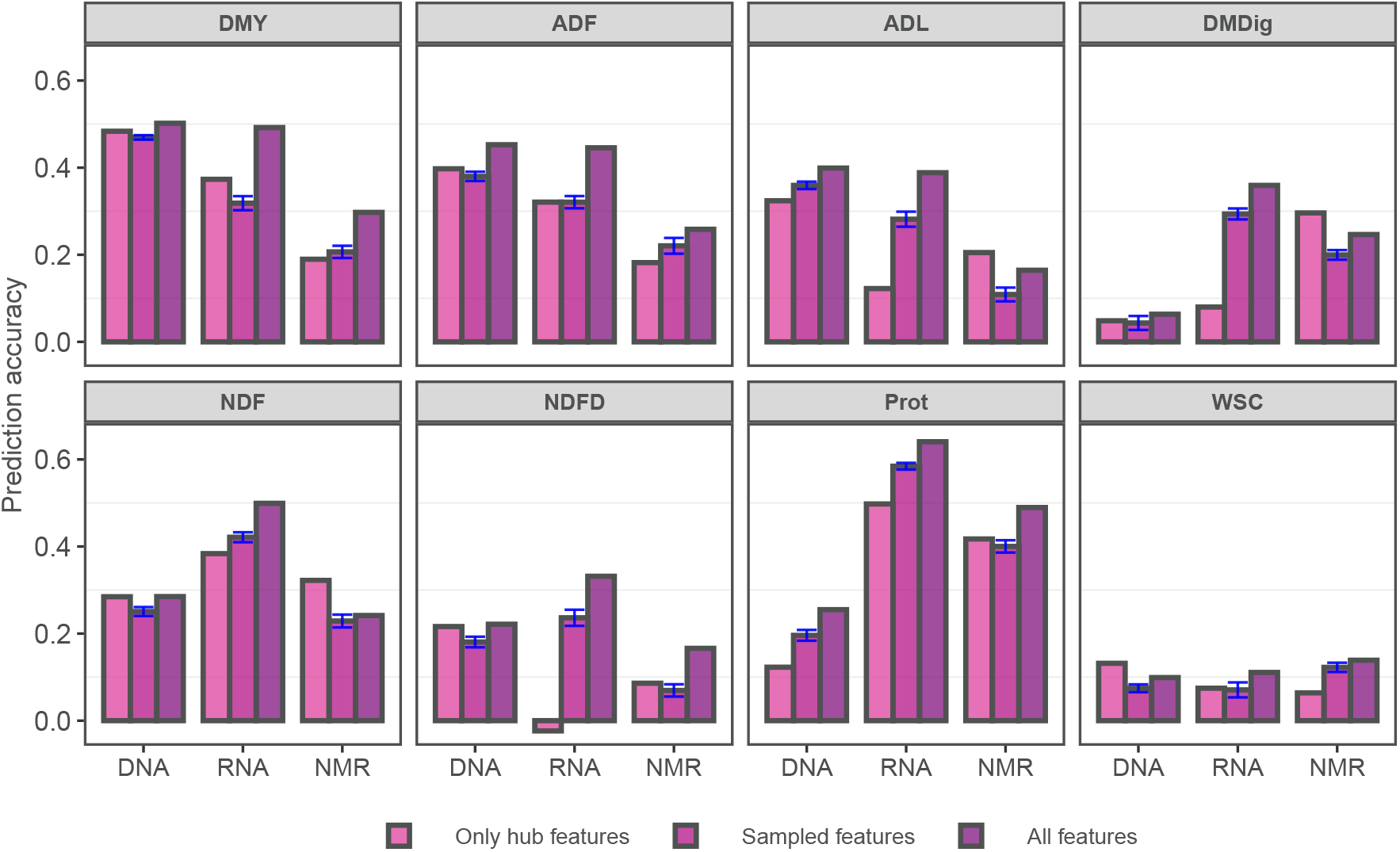
Random forest-based prediction accuracy computed for eight forage grass traits as a function of predictors encompassing three omic layers (DNA: SNP-based markers, RNA: gene expression via RNA-seq, and NMR: variables representing bucketed NMR spectra) and three predictor set configurations as indicated by the color gradient. The standard errors for the mean accuracy of sampled features are depicted in blue color. DMY - dry matter yield; ADF - acid detergent fiber; ADL - acid detergent lignin; DMDig - digestible dry matter; NDF: -neutral detergent fiber; NDFD - digestible NDF; Prot - protein; and WSC - water-soluble carbohydrates.

## Discussion

The study elaborated here explores a network-based approach to combine multi-omic data arising from an *n* ≪ *p* scenario, inferring associations between biomarker candidates with dry matter yield and nutritive quality traits of polyploid forage grass families. This was accomplished by using a joint graphical lasso model with a fused penalty for network reconstruction, followed by topological property extraction and integration via multivariate mixed modeling. Further, a machine learning-based prediction scheme was explored to verify the extent of information available in hubs and in the whole feature space for predicting agronomically important phenotypes. The plant material consisted of family pools of inter-specific and-generic grass hybrids from two connected diallels. Crossing different pasture species/genera is not a trivial task; obstacles can emerge. Firstly, out of all initially planned crosses, only a subset generated viable seeds, impacting the sample size. Also, seeds were not abundant for many of the crosses, requiring an additional year of multiplication. Secondly, extraneous offspring patterns were detected, prompting a question of whether normal parental contributions were formed for some of the F_2_ families. This inquiry remained unanswered in this manuscript given the complexity of the genetic material (family pools), SNPs called from RNA-seq data, and the unavailability of parental genotypes. Despite the self-incompatibility (SI) ensuring cross-pollination in perennial ryegrass (Cropano et al. 2021), four to eight percent of self-fertilization has been reported (Arcioni and Mariotti 1983;Deniz and Dogru 2007). This, in addition to the low success rate of inter-specific and-generic hybridization, might have caused the deviated genomic state of offspring families for crosses that produced a small number of seeds. We did not use the parental information from the diallel structure in the network construction but removed it by regression to control for the kinship among individuals across analyses, a crucial action to avoid spurious results in network reconstruction. Due to the genetic design, correlation among samples is expected, which can lead to the detection of co-expression among features as a result of shared chromosomal segments. Additionally, confounding artifacts not controlled for can affect groups of genes and NMR variables, which can lead to the detection of spurious correlations. We fitted population structure as covariates by using principal component scores derived from the genetic markers covering the whole genome aiming to alleviate the non-independence among samples, which has been shown to reduce false network discoveries efficiently (Parsana et al. 2019). An extra layer of precaution to avoid the effect of false-positive edges was deployed by retaining only common hub features between pedigree classes.

The gene co-expression and metabolic networks as the ones we reconstructed in this study (Figure 4) using RNA-seq and NMR variables, respectively, can contain interesting topological properties e.g., the existence of highly connected nodes and the organization of nodes in modules (Li et al. 2015). We explored these two properties aiming to select, across pedigree classes, conserved hubs extracted at a rate of five per module, therefore, increasing the likelihood of sampling hubs associated with diverse biological processes. Our approach to selecting and associating these features with phenotypic traits is altogether different from the conventional method, which consists of performing a simple correlation-based gene co-expression network analysis followed by thresholding to find modules that can then be summarized into a synthetic (eigen) gene for association with external sample traits (Langfelder and Horvath 2008). As highlighted by other authors(Huynh-Thu and Sanguinetti 2018;Jiang et al. 2019), this correlation-based approach cannot distinguish between linear relationships due to directly dependent nodes and those arising from confounding nodes, which might create spurious edges in the graph and, consequently, misleading clustering. In contrast, Gaussian graphical models, as used here, are based on the precision (inverse variance) matrix and express conditional dependence between pairs of features given all the other variables in the data set (Danaher et al. 2014) which, therefore, avoids declaring an edge when no causal relationship exists. Regarding the presence and distribution of edges across reconstructed networks, the proportion of undirected edges given the total available nodes was much higher for the NMR -based metabolic network relative to the gene expression graph. This is a consequence of the lack of independence among bins closely located across the NMR spectrum. Indeed, an average autocorrelation across samples revealed significant spikes up to lag 12 (data not shown). Therefore, a proper feature selection algorithm for spectral data can be implemented to deal with the existence of autocorrelation.

Picturing a biological regulatory cascade, hub genes are usually regulatory factors located upstream, whereas genes represented by low-degree nodes are located on the other end(Zeng et al. 2022). They can be associated with biological processes from which several others are dependent, yielding the commonly observed power-law degree distribution. The presence of a limited amount of important hub genes, however, does not necessarily imply a simple genetic architecture, because the regulation of the hub gene expression is typically highly polygenic. Investigating putative hubs can reveal important genes as, for example, the cold-regulated gene *Lolium perenne LIR1* (*LpLIR1*) (Ciannamea et al. 2007) represented by the hub gene coded as 22 in Figure 5, which is located at chr5:155166187-155167265 in the *L. perenne* genome and appears to act in the photoperiodic regulation of flowering. Another example is hub 7, which represents the *PDX1.1* gene, involved in the biosynthesis of vitamin B6 and protection against stresses (Liu et al. 2022). Overexpression of PDX proteins was shown to increase seed size and biomass in Arabidopsis (Raschke et al. 2011). For metabolite-metabolite networks, high-degree nodes may represent signaling molecules or molecules engaged in many reactions. The content and diversity of such molecules have been shown to be shaped by domestication as well as due to crop improvement (Alseekh et al. 2021). Improving biomass output per area is the ultimate breeding goal in a forage breeding program and also implies selection pressure for stress endurance due to animal grazing or mechanical harvesting. In this sense, secondary metabolites are well-known for their role in the plant’s response to external disturbances as herbivory (Degenhardt Jorg 2009). In more general, significant associations can be detected between metabolites and agronomic traits (Turner et al. 2016) and the whole NMR spectrum can be used for metabolomic-assisted prediction (Guo et al. 2022). That being stated, genetic selection for elite grasses might be linked to an altered profile of metabolites, leveraging their usefulness as markers for selection or for prediction purposes. Indeed, great chemical diversity is available in perennial ryegrass (Subbaraj et al. 2019), not only adding another layer of information for omics-assisted breeding but also enabling target improvement of varieties with a specific profile of key metabolites.

Together, significant additive genetic correlations between omic features and phenotypic traits displayed in Figure 5 and the presence of over-represented gene ontology (GO) terms in the hub gene set supports the evidence that these features hold fundamental biological properties. We further assessed the predictive power available in the sets of gene and metabolite hubs. This was accomplished by merging the HR and FL data sets for trait prediction aiming to increase the sample size, which even though still below the appropriate size for genomic selection was counterbalanced by a high signal-to-noise ratio given the diallel structure which is expected to boost information for model learning (see Figure 1). Splitting between training and testing sets would reduce the sample size for training. Therefore, we used the ensemble learning method of random forest with all samples and reported the out-of-bag (OOB) accuracy as a prediction performance metric, eliminating the need to set aside a test set(Breiman 2001). Despite the crossing scheme, eigenvectors from marker data did not reveal large dissimilarity between pedigree classes (Supplemental Figure S1), therefore allowing for the joint analysis. Also, random forest is not very sensitive to hyperparameter tunning (Probst et al. 2019), making it a good option for the designed prediction setup. This can be attested by the magnitude of predictions displayed in Figure 6. Prediction accuracy for dry matter yield was reported in other studies at 0.31 using diploid ryegrass synthetic populations (Pembleton et al. 2018), 0.34 using tetraploid ryegrass (Guo et al. 2018), and 0.5 investigating diploid perennial ryegrass(Arojju et al. 2020). Here, we report values of prediction accuracy of dry matter yield that approximate 0.5 (Figure 6) using both SNP-markers and gene expression, despite the lower sample size but helped by high relatedness among samples, an important component in genomic selection (Edwards et al. 2019). Also for dry matter yield, surprisingly the most heritable trait (Supplemental Table S1), the set of hub genes and SNPs markers tagging them seem more predictive than features sampled at random. For the remaining traits, mixed results were observed which can be an artifact due to sample size, low heritability, or population structure. Additionally, the signal might be dependent on the genetic background and disappeared as we merged the two data sets for the prediction study. Heritability is an important parameter driving prediction accuracy. If it is low, the error variance will be higher, leading to difficulties in estimating the effect of genome segments accurately (van der Werf 2013), especially if the sample size is not sufficiently large. Small values of heritability were primarily observed for quality traits (supplemental Table S1), which explains the lack of predictive power of the model for digestibility, water-soluble carbohydrates, and digestible NDF, for example. The NIR-based quality parameters are obtained from calibrated models using data of chemical analysis from samples of standard breeding materials and might not translate well into curves of inter-generic and-species hybrids, explaining the lower heritability.

Given that plant tissues were sampled once from pools of seedlings grown in a greenhouse environment at the F_2_ generation for transcriptomic and metabolomic analyses, the information carried by the recorded features represents a snapshot of the complex interactome at that particular condition in space, time, and random mating generation. This information was learned by the model and translated into higher prediction accuracy for protein and digestibility, despite the fact that phenotypes were recorded in later growth stages and in another generation of random mating. Across omic layers, the results also showed that using all available features is almost always a better choice for increased prediction accuracy. Besides more main effects being captured, the random forest model can capture feature-feature interactions (Yao et al. 2013) as long as the marginal effects are large enough to cause a tree split, therefore, accounting for some of the existing epistasis. Therefore, the existence of significant edges displayed in Figure 5 and the magnitude of the prediction accuracies presented in Figure 6 reveals a strong link between field-based phenotypes and heritable omic features assessed from young seedlings in a controlled environment. Altogether, this information brings the question of whether phenotypes from seedlings grown for DNA sampling could be recorded through a low-cost NIR-based method and used to improve the accuracy of genomic selection models, a subject worthy of consideration in future research.

The use of multi-omics in plant breeding-related studies is becoming more popular due to decreasing in cost per data point as a result of modern high-throughput technologies. This has been allowing researchers to reconstruct complex biological networks for inference and mining. Out of the many topological properties that can be retrieved from an interaction network, hub features showing many putative links have been shown to play important biological roles in plants (Tahmasebi et al. 2019). Our study reveals that narrowing down the high-dimensional feature space generated by high-throughput omic analysis to fewer entities by leveraging properties of the graphical theory can reveal important biomolecules for molecular studies and breeding. Additionally, dimensionality reduction can substantially boost detection power by alleviating the multiple testing problem. Further investigations of candidate features may help elucidate biological processes underlying the expression of phenotypic traits and serve as markers for omics-assisted selection in breeding programs. Even though we did not perform compound identification from the NMR data, this is a feasible task and may reveal metabolites playing important roles in biomass yield and nutritional quality.

## Conclusion

The scientific community has seen a sharp increase in publications exploring the usefulness of biological network reconstruction based on high throughput omics data since the 2000s, but studies with forage species remain scarce. Here, we have explored the usefulness of topological properties of gene co-expression and metabolic networks in explaining the phenotypic variance of eight traits assessed in family pools of inter-specific and -generic grass hybrids. Network topology estimated via fused graphical lasso revealed profound network differences between pedigree classes, but a set of 30 high-degree hub genes and 32 hub NMR variables remained conserved across classes given the selection criteria, out of which 10 hubs were found as candidate biomolecules significantly associated with the expression of agronomic phenotypes. Gene set enrichment analysis and weighted omics-phenotype network estimation suggested that sets of hubs are likely to contain essential features modulating interactomes and the expression of economically important phenotypes.

## Abbreviations

ADF: acid detergent fiber
ADL: acid deterged lignin
BIC: Bayesian information criterion
BLUE: best linear unbiased estimator
DMDig: dry matter digestibility
DMY: dry matter yield
FDR: false discovery rate
FL: *Festulolium loliaceum*
GBS: genotyping-by-sequence
GO: gene ontology
GRM: genomic relationship matrix
HR: hybrid ryegrass
IQR: interquartile range
JGL: joint graphical lasso
LRT: log-likelihood ratio test
MAF: minor allele frequency
NDF: neutral deterged fiber
NDFD: digestible NDF
NIR: near-infrared spectroscopy
NMR: nuclear magnetic resonance
OOB: out-of-the-bag accuracy
PC: principal component
Prot: protein
REML: restricted maximum likelihood
RNA-seq: RNA sequencing
SNP: single nucleotide polymorphism
WSC: water-soluble carbohydrate

## Acknowledgments

This work was supported by the Innovation Fund Denmark, through the project Breed4Biomass (grant 6150-00020B). The authors gratefully acknowledge the financial support provided by the aforementioned organization. The authors declare no competing interests.

## Author contribution statement

EB conceptualized the study, wrote the code for analyses and data visualizations, interpreted the results, and drafted the manuscript. DF contributed to the conception and design of the B4B project and data curation. IN contributed to omics data generation and miscellaneous bioinformatics analyses. IL contributed with bioinformatics expertise and project design. MG created the populations, carried out field trials, and recorded agronomic phenotypes. TD converted NIR spectrums into nutritive quality parameters. CSJ, TA, and LJ contributed to the conception and design of the B4B project and funding acquisition. TA acquired funding for and coordinated the B4B project. LJ supervised the current study and provided valuable comments. All authors critically reviewed the manuscript. All authors have read and approved the manuscript.

## Data availability statement

The omics data sets supporting the conclusions of this article have been made available through an R data package named “breed4biomass”, which is openly available on GitHub (https://github.com/elesandrobornhofen/breed4biomass).

## Supplemental Material

### Supplemental figures

**Figure S1:**
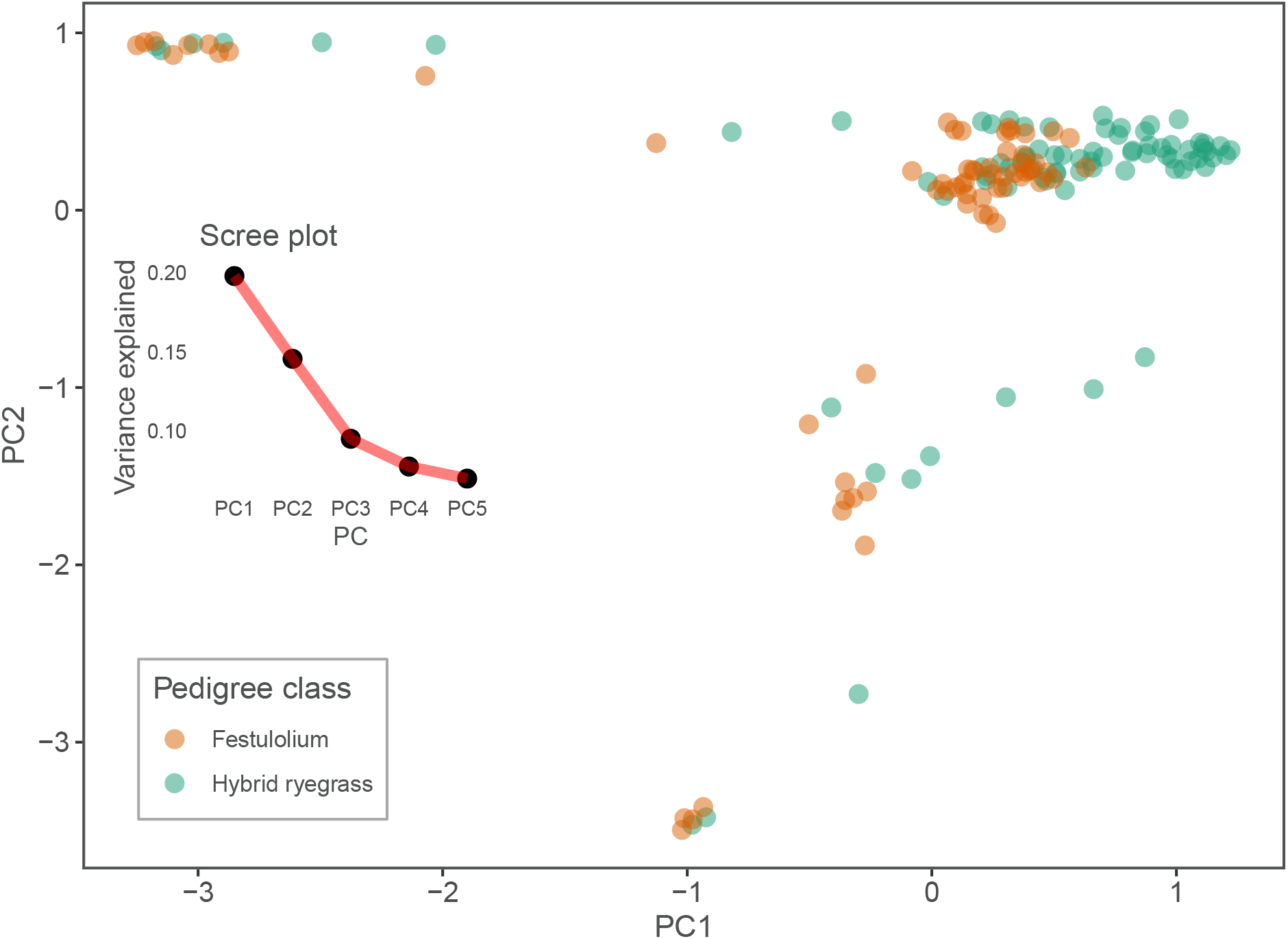
Scatter plot displaying scores of the first two principal components (PCs) from the PC analysis of the combined (hybrid ryegrass plus *Festulolium loliaceum* samples) genomic relationship matrix. The number of samples is equal to 144. An overlaying scree plot shows the variance explained by the first five PCs.

### Supplemental tables

**Table S1:** Restricted maximum likelihood (REML) estimation of variance components from two field trials comprising different pedigree classes of forage grasses: HR - hybrid ryegrass and FL - *Festulolium loliaceum*.

| Ped.<br>class | Variance<br>component | Traits<sup>1</sup> |  |  |  |  |  |  |  |
| --- | --- | --- | --- | --- | --- | --- | --- | --- | --- |
|  |  | DMY | ADF | ADL | DMDig | NDF | NDFD | Prot | WSC |
| HR | $\sigma_u^2$ | 2181.088 | 0.125 | 0.002 | 0.135 | 0.278 | 1.451 | 0.186 | 0.000 |
|  | S.E. | 1054.606 | 0.073 | 0.002 | 0.086 | 0.174 | 0.741 | 0.113 | 0.368 |
| FL | $\sigma_u^2$ | 7353.689 | 0.260 | 0.018 | 0.148 | 0.590 | 0.000 | 0.034 | 0.194 |
|  | S.E. | 3090.586 | 0.128 | 0.011 | 0.281 | 0.277 | 0.540 | 0.046 | 0.162 |
| HR | $\sigma_f^2$ | 0.000 | 0.000 | 0.000 | 0.031 | 0.029 | 0.000 | 0.000 | 0.269 |
|  | S.E. | 892.753 | 0.075 | 0.002 | 0.099 | 0.183 | 0.859 | 0.117 | 0.587 |
| FL | $\sigma_f^2$ | 0.000 | 0.000 | 0.000 | 0.374 | 0.102 | 0.704 | 0.071 | 0.000 |
|  | S.E. | 1760.040 | 0.099 | 0.008 | 0.441 | 0.186 | 0.976 | 0.066 | 0.200 |
| HR | $\sigma_s^2$ | 183.365 | 0.206 | 0.007 | 0.222 | 0.527 | 0.253 | 0.377 | 1.530 |
|  | S.E. | 721.615 | 0.106 | 0.003 | 0.136 | 0.263 | 0.547 | 0.193 | 0.810 |
| FL | $\sigma_s^2$ | 4431.405 | 0.443 | 0.000 | 0.000 | 0.402 | 0.228 | 0.039 | 1.835 |
|  | S.E. | 2338.886 | 0.194 | 0.008 | 0.443 | 0.254 | 0.927 | 0.056 | 0.599 |
| HR | $\sigma_e^2$ | 4821.160 | 0.388 | 0.010 | 0.512 | 0.925 | 4.189 | 0.561 | 3.242 |
|  | S.E. | 986.994 | 0.085 | 0.002 | 0.115 | 0.210 | 0.722 | 0.139 | 0.740 |
| FL | $\sigma_e^2$ | 5939.760 | 0.361 | 0.046 | 2.400 | 0.728 | 5.799 | 0.319 | 0.659 |
|  | S.E. | 1618.810 | 0.114 | 0.010 | 0.528 | 0.202 | 1.225 | 0.073 | 0.286 |
| HR | $\mu$ | 1312.640 | 23.420 | 2.110 | 88.937 | 42.872 | 71.172 | 14.143 | 10.986 |
|  | S.E. | 20.718 | 0.301 | 0.051 | 0.326 | 0.477 | 0.626 | 0.391 | 0.824 |
| HR | $\mu$ | 1204.610 | 26.757 | 2.912 | 85.493 | 46.501 | 70.291 | 11.720 | 12.422 |
|  | S.E. | 50.153 | 0.442 | 0.084 | 0.603 | 0.517 | 0.942 | 0.255 | 0.782 |
| HR | $h^2$ | 0.304 | 0.174 | 0.126 | 0.150 | 0.158 | 0.246 | 0.165 | 0.000 |
| FL | $h^2$ | 0.415 | 0.244 | 0.288 | 0.051 | 0.324 | 0.000 | 0.074 | 0.072 |

<sup>1</sup>DMY: dry matter yield; ADF: acid detergent fiber; ADL: acid detergent lignin; DMDig: digestible dry matter; NDF: neutral detergent fiber; NDFD: digestible NDF; Prot: protein; and WSC: water-soluble carbohydrate. $\sigma_u^2$ , $\sigma_f^2$ , $\sigma_s^2$ , and $\sigma_e^2$ are the genomic variance, variance due to uncorrelated family effects, spatial variance, and residual variance, respectively. S.E. is the asymptotic standard error and $h^2$ is the genomic heritability.

## Notes

### Competing Interest Statement

The authors have declared no competing interest.

